# Correlated dipolar and dihedral fluctuations in a protein

**DOI:** 10.1101/2021.01.28.428566

**Authors:** Abhik Ghosh Moulick, J. Chakrabarti

**Author notes:** Also at Thematic Unit of Excellence on Computational Materials Science and Technical Research Centre S. N. Bose National Centre for Basic Sciences, JD Block, Sector – III, Salt lake, Kolkata-700106, India.

## Abstract

Correlation between dihedral fluctuations is a possible way to understand coordination between various amino acid residues of the protein. The nanosecond timescales of correlated fluctuations of dihedral angle do not allow direct probing by experimental methods. However, NMR experiments probe dipolar fluctuations given in terms of cross correlated relaxation (CCR) rates, expressed as zero frequency spectral density function (J) of the fluctuations of the mutual orientation of two spatially separated dipole vectors. Here we illustrate the correlation of protein dipolar and dihedral angle using molecular dynamics simulation of protein Ubiquitin (Ub) and GB3. We calculate CCR rates between protein bond vector from simulation and compare with CCR data obtained from NMR experiments. A good correlation between theoretical and experimental values is found. We further show that the zero frequency spectral functions of backbone dihedral *ψ* auto-correlation function and dipole orientation fluctuations show strong correlations. These correlations are not sensitive to protein and forcefield parameters. Hence, CCR may act as a marker for protein backbone dihedral fluctuations.

## I. INTRODUCTION

Ligand binding sites in a protein are often separated by distances much larger than atomic sizes^1–4^. However, they show correlated behaviour while binding to ligands. This correlation is usually expressed in terms of the Pearson correlation coefficient^5–10^. The Pearson correlation coefficient, given by covariances of two variables, is a purely static quantity. In the case of functional residues, the correlations have intrinsic timescales that control the rate of binding events. It has been shown that one can generalize the covariance between two variables with a time lag to construct time dependent correlation function (TDCF)^11^ of fluctuations of two dihedral angles of functionally relevant residues^12^. The time dependent correlations of dihedral fluctuations persist till nanosecond (ns) timescales ^12^. Although the dihedral fluctuations are theoretically useful, the nanometer timescales of the correlated fluctuations down to the spatial resolution of the residues are not yet directly amenable to experimental probes. It is interesting to ask if the dihedral fluctuations can be connected to an experimentally measurable quantity that probes microscopic motions in proteins.

Nuclear magnetic resonance (NMR) techniques^13^ are used to probe atomic motions through dipolar fluctuations in a protein^14–37^. The collective dynamics of a group of atoms^38–40^ are given in terms of cross correlated relaxation (CCR)^41^ rates, which is the zero frequency spectral density function of the fluctuations of the mutual orientation of two spatially separated dipole vectors^42,43^. Experiments often consider fluctuations in mutual orientation between *H^N^* - *N* and *C_α_* - *H_α_* dipole pairs. Such data have been reported for GB3^44^. The representative dipole pair *H^N^* - *N/C_α_* - *H_α_* is shown in Fig. 1(a). The backbone dihedrals are also shown in the same figure. The angle between 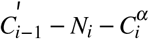 and 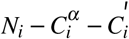 planes, illustrated in the figure, is the backbone dihedral *ϕ*. Similarly, the angle between 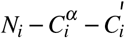 and 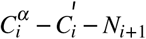 planes is the backbone dihedral *ψ*. These backbone dihedrals involve the dipole pairs 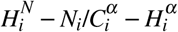. Hence, one would expect the backbone dihedral to be sensitive to the fluctuations of mutual orientation of these dipoles. In an earlier work^45^ we show correlated nature between *H^N^* - *N/C_β_* - *H_β_* dipole pair and dihedral angle *ψ* fluctuations for some residues of Ub. While the dihedral fluctuations take place in nanoseconds, the CCR rates provide integrated information over time on dipole orientation correlation function. If the dipolar fluctuations at timescales of nanoseconds have an important contribution to CCR, CCR may act as markers for backbone dihedral fluctuations as well. Motivated by this, we examine the zero frequency spectral functions of intra-residual backbone dihedral angles and *H^N^* - *N/C_α_* - *H_α_* dipolar orientation fluctuations in two proteins GB3 and Ubiquitin (Ub).

**FIG. 1.**
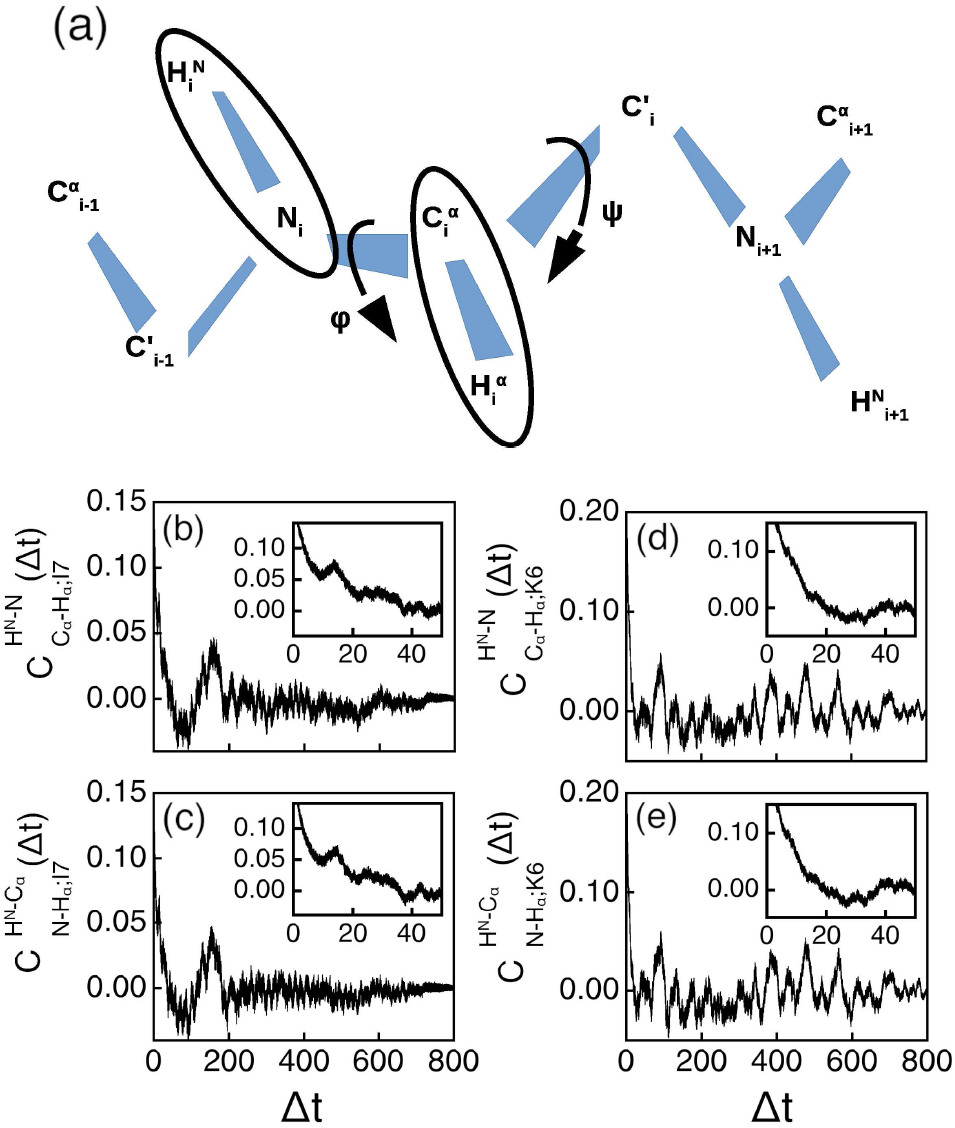
(a) Cartoon representation of dipoles (within ellipses) used in the present study. The backbone dihedral angles *ϕ* and *ψ* are shown by arrows. TDCFs of dipolar orientational fluctuations for Isoleucine, I7 of GB3 (b) 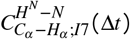, and (c) 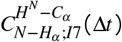. Similar TDCFs for Lysine, K6 (d) 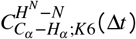 and (e) 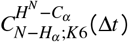, of Ub. Insets show short time behaviour of the correlation functions.

We use long trajectories of the proteins in molecular dynamics (MD) computer simulation to compute the time dependent correlation functions over the equilibrated part of the trajectory. The time dependent correlation functions are constructed as follows: The dipolar fluctuations for an angle between dipole pair *H^N^* - *N/C_α_* - *H_α_*, while the backbone dihedral angles *ϕ* and *ψ* of the given residue. Typical correlations persist up to tens of nanoseconds. We extract the zero frequency spectral functions by numerically integrating the correlation functions over this timescale. We find that the experimental data for GB3 compare well with our theoretical CCR data for dipolar orientation fluctuations. Thus the experimental CCR is largely accounted for by those obtained by restricting the integration time up to a few tens of nanoseconds. This indicates that the nanosecond timescale dipolar fluctuations dominate the experimental CCR. We find good correlations between zero frequency spectral density of functions involving backbone dihedral *ψ* and orientational fluctuations of dipoles *H^N^* - *N/C_α_* - *H_α_* for both the proteins.

## II. METHODS

### A. SIMULATION DETAILS

We consider GB3 (PDB id: 2OED)^46^ with 56 amino acids and ubiquitin (PDB id: 1UBQ)^47^ with 76 amino acids in our studies. Amber99sb force field (ff)^48^ is used in the GRO-MACS software package^49^ for simulation. The TIP3P water model is considered for solvent molecules and counterions are added for electroneutrality. Particle Mesh Ewald method is used to assess long ranged electrostatic energy^50^. 10 Angstrom (Å) is considered as truncation limit for both Lennard-Jones and short range interactions. GB3 protein is solvated in a cubic box of dimensions 5.6×5.6×5.6 nm with 5524 water molecules 2 Na^+^ ions are added to attain electrical neutralization. Ubiquitin is solvated in a cubic shaped box of dimensions 6.49×6.49×6.49 nm^3^ with 8696 water molecules. No counter ions are added to Ubiquitin as the system is already neutral. Minimization is done for 50,000 steps using the steepest descent algorithms. All bonds in the protein are constrained using the LINCS method. Equations of motion are integrated using leap-frog algorithm with an integration time step of 2fs. Systems are equilibrated through 2 steps (NVT & NPT) using position restraints to heavy atoms. NVT and NPT equilibration is carried out at 300K Temperature and 1 Bar pressure. Simulations are executed for 1.05 *μs* with 2 femtosecond time step integration employing periodic boundary conditions in all directions. In the case of Ub, we further perform molecular dynamics simulation using the CHARMM27^51^ force field in the NAMD^52^ software, following standard protocols for the NPT ensemble. All other parameters are kept the same as in the GROMACS simulations. The calculations of root-mean-square deviation are done with respect to the backbone atoms. Visual analysis of protein structure is done using the Pymol.

### B. ANALYSIS

Time dependent correlation function (TDCF) between two dynamic variables A and B for time interval t is given by,

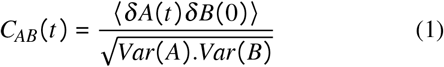

where *δA*(*t*) = *A*(*t*) - *Ā* and 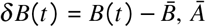 and 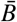 the mean values, and Var(A) and Var(B) the variances of A and B. Angular brackets signify average over initial times. When *A* = *B*, the corresponding correlation is called autocorrelation function, whereas *A* ≠ *B* corresponds to crosscorrelation function. We compute the autocorrelation function of temporal fluctuations of the cosine of the angle between *H^N^* - *N* and *C_α_* - *H_α_* dipole pairs in Eq.1. We also compute the TDCF for dihedral fluctuations using sine of the dihedral angles as dynamical variables using Eq.1.

TDCF for the dihedral angle from simulation trajectory has been computed as follows: For any given time difference *Δt* = |*t*_2_ -*t*_1_|, the product is computed between fluctuations of dihedral *θ* and *θ’* of residue i for the l-th observation for a given *Δt*,

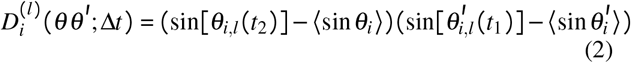

Here angular brackets denote ensemble average of the corresponding quantity over the entire simulation trajectory. The TDCF is given by:

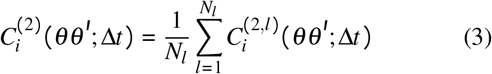

where,

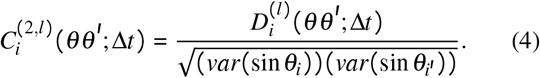

Here *N_l_* corresponds to the number of observations for a given *Δt*. When *θ = θ’* corresponding correlation is called autocorrelation. We compute the TDCF for dipolar fluctuations similarly.

The spectral density function *J*(*ω*) is defined as Fourier transform of the correlation function *C*(*t*)^53,54^.

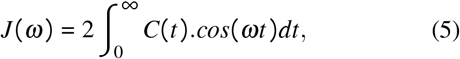

which corresponds to an integral over time for zero frequency. The integrals are computed numerically from the smoothened autocorrelation functions using the Gaussian quadrature^55^. The data sets of time dependent correlation functions are smoothened using cubic spline fit technique^56^. The smoothening spline for a function f(x) minimizes 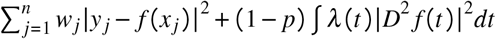, where the first term measures error and the second term represents roughness measure. Error measure weights *w_j_* are equal to 1 by default. *D^2^ f* denotes the second derivative of function f. The smoothing parameter *p* is in the range [0,1]. Here smoothening is done using *p*=0.3.

## III. RESULTS AND DISCUSSIONS

The equilibration of the all-atom MD simulations of the system is ensured from the saturation of root-mean-square deviation (RMSD) of the backbone atoms, as shown in Supplementary information (SI) Fig.S1 for protein GB3. SI Fig.S2 shows RMSD plot for Ub. We perform the analysis over the equilibrated part of the trajectory. To validate the simulations we compare the crystal structure of the proteins with the average structures over the equilibrated trajectory. The aligned conformations for both structures are in SI Fig.S3 for GB3 and SI Fig.S4 for Ub. For GB3, all heavy-atom RMSD is 1.21 Å and for Ub, RMSD is 1.80 Å. We also compare *ψ - ϕ* Ramachandran plot for both the crystal and average structures for both proteins SI Fig.S5 - Fig.S6. The Ramachandran plots show good structural similarities of both the proteins with the experimental crystal structures. To avoid artefact due to the rigid rotation, we superimpose all conformations with respect to the reference structure chosen for both proteins at t = 250 ns after which the RMSDs for both are saturated and calculate the angles from the superimposed structures. We construct the TDCF for all the relevant variables and compute the zero frequency spectral functions by numerically integrating smoothened correlation function data.

*TDCF:* First, we consider the TDCF of fluctuations of dipolar orientation. We consider the dipole pairs *H^N^* - *N* and *C_α_ - H_α_*. Let *θ* be the angle between a pair of dipoles *H^N^* - *N* and *C_α_ - H_α_* of a residue *R*. The values of *P_2_*(cos *θ*) describes the dipolar orientation fluctuations. Experimental CCR data cannot distinguish between the orientation fluctuations between the direct and the cross pairs, *H^N^ - N/C_α_ - H_α_* and *H^N^ - C_α_/N - H_α_*. Hence, we consider both the pairs in our calculations. Let us illustrate the case of Isoleucine (I7) of GB3 which belongs to the beta-sheet of the protein crystal structure. Figs. 1(b) and (c) show the TDCFs 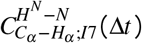 and 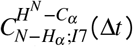, respectively. The correlation functions decay within 40 ns, along with oscillations of small amplitudes about zero at larger times. The short time behaviors are illustrated in the insets for both the cases. The temporal correlation functions for the other cases show similar behaviour. We also show typical cases of dipolar fluctuations of Ub. Fig. 1(d) and (e) show 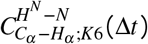 and 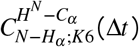 respectively, for Lysine, K6. This residue belongs to the beta sheet region in the crystal structure. Here again, the TDCFs decay within 20 ns to zero value (Fig. 1(d) and (e)). The short time decays in both cases are shown in the respective insets. The dipolar orientation fluctuations of the other residues show similar behaviour. Some cases have been illustrated in SI Figs (Fig.S7).

We denote the TDCF of dihedral fluctuations by *C_R_*(*θθ’,Δt*) where *θ* and *θ’* are the backbone dihedral of residue R. We calculate the dihedral angles for backbone *ϕ* and *ψ* from the atomic positions, the sine of these functions being used as dynamical variables for the dihedral fluctuations. We show backbone dihedral fluctuations for representative cases in Fig. 2 where the insets show the short time data. Figs. 2(a)-(c) show cases for GB3. *C_I7_*(*ϕϕ,Δt*) decays to zero in 30 ns (Fig. 2(a)). *C_I7_*(*ϕψ,Δt*) starts with anticorrelation at *Δt* = 0 and decays to zero within 40 ns (Fig. 2(b)). *C_I7_*(*ψψ,Δt*) decays within 30 ns (Fig. 2c). All three cases reveal that at long times the TDCFs fluctuate around the zero value (Figs. 2a-c). As representative cases of TDCF for Ub, we show the dihedral correlation functions for Lysine, K6 in Figs. 2(d)-(f) and the small time behaviour in the corresponding insets. *C_K6_*(*ϕϕ,Δt*) loses initial correlation within 20 ns (Fig 2d and inset). *C_K6_*(*ϕψ,Δt*) (Fig. 2(e)) is initially anticorrelated at *Δt* = 0, and the initial anti-correlation decays within 20 ns (inset Fig. 2e). *C_K6_*(*ψψ,Δt*) fluctuates around zero value following initial decay within 20 ns (Fig. 2(f), and inset). We find similar decay of correlations in other cases as shown for few more cases for both the proteins in SI Fig.S8. The overall nature of TDCF of the dihedral fluctuations is in agreement with the previous report ^12^.

**FIG. 2.**
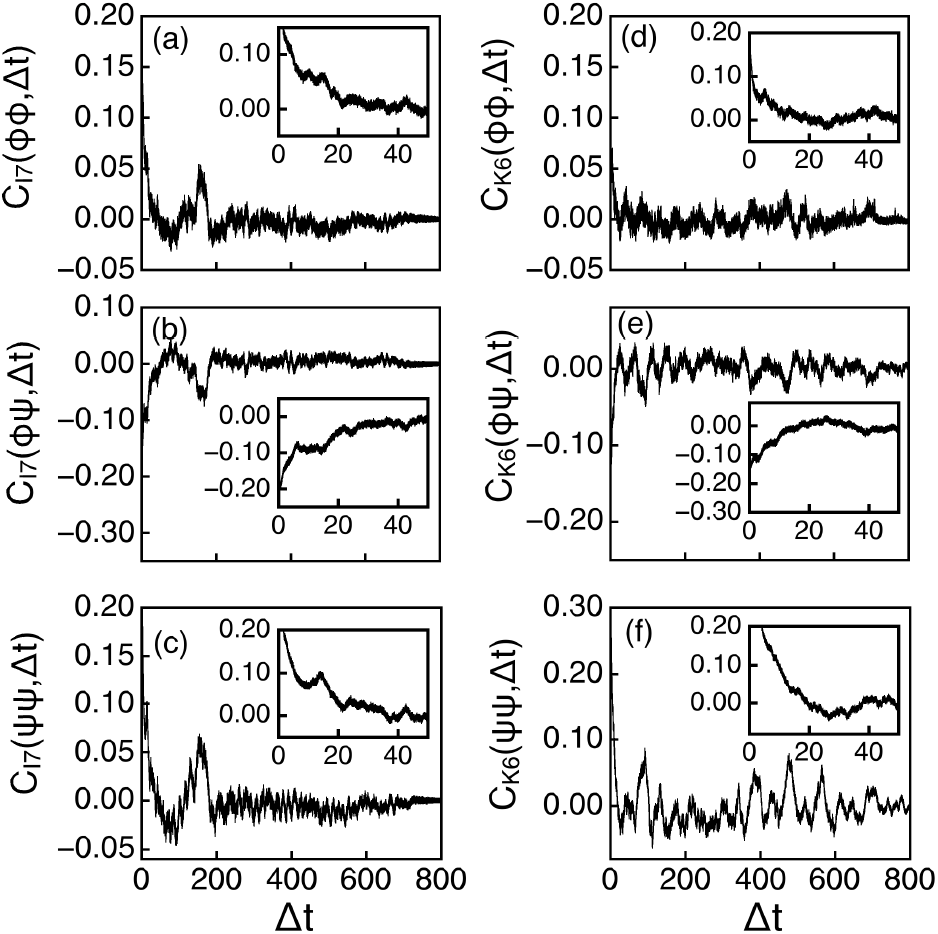
TDCFs between various dihedral fluctuations of Isoleucine, I7 of GB3:(a) *C_I7_* (*ϕ ϕ, Δt*), (b) *C_I7_* (*ϕ ψ, Δt*), and (c) *C_I7_* (*ψψ, Δt*); (d) *C_K6_* (*ϕϕ, Δt*), (e) *C_K6_* (*ϕψ, Δt*), and (f) *C_K6_* (*ψψ, Δt*) of Lysine, K6 of Ub. Insets show short time behaviour of the TDCFs.

We examine the timescales of decay of the correlation functions. Since the long time oscillations have low amplitude, we restrict only to the short time regime shown in the insets of Fig.2 to extract the decay timescales. We fit the data to an exponential form, 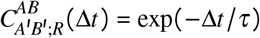 where *τ* is the correlation timescale. The timescale for dipolar orientation is denoted by *τ_R, H^N^-N/C_α_-H_α__* for *H^N^* - *N/C_α_ - H_α_* dipole pair and *τ_R,H^N^-C_α_/N-H_α__* for *H^N^* - *C_α_/N - H_α_* cross-dipole pair. The correlation timescale for dihedral fluctuations is calculated in a similar fashion. The timescale for *ϕϕ* fluctuations are denoted by *τ_R,ϕϕ_*, those for *ϕψ* correlations by *τ_R,ϕψ_* and for *ψψ* fluctuations by *τ_R,ψψ_*. In the case of anti-correlations, we consider the modulus of the values for fitting. Fig. 3(a) shows the histogram *H^dipole^* Of *τ_R,H^N^-N/C_α_-H_α__* and *τ_R,H^N^-C_α_/-N-H_α__* considering dipolar fluctuations of all the residues of both proteins. The combined correlation timescale is denoted by *τ^dipole^*. The histogram shows that correlation timescale extends to about 10 ns. Fig. 3(b) shows histograms *H^dihedral^* considering the timescales of the dihedral autocorrelations *τ_R,ϕϕ_* and *τ_R,ψψ_* and cross-correlations *τ_R,ϕψ_*, taking care of all the residues of the two proteins together. We denote the combined correlation times simply by *τ^dihedral^*. The histograms show that the timescales of correlations for both the dipolar mutual orientation and dihedral fluctuations mostly lie within 20-30 ns. This observation is in agreement with earlier work that dihedral fluctuations occur in nanosecond timescale^12^.

**FIG. 3.**
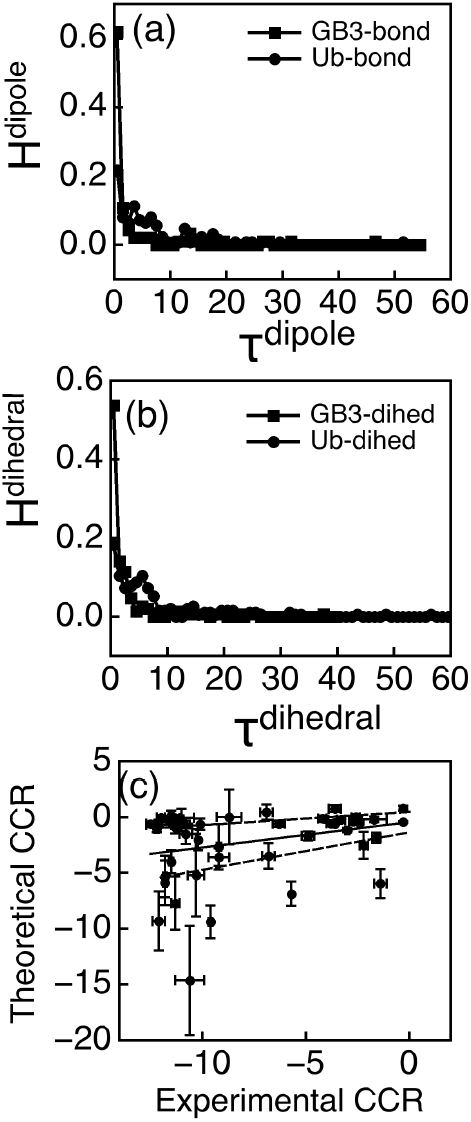
(a) Histogram (*H^dipole^*) of autocorrelation timescale for both GB3 and Ub considering dipole pair (a) *H^N^* - *N/C_a_* - *H_a_* (both direct and cross pairs). (b) Histogram (*H^dihedral^*) of correlation timescales for dihedral angle fluctuations *τ^dihedral^* which consist of *τ_R,ϕϕ_*, *τ_R,ϕψ_* and *τ_R,ψψ_* for both proteins. (c) Correlation diagram between CCR rates of experimental and theoretical values for GB3 along with the error bars.

*Zero frequency spectral functions and CCR:* We check if the fluctuations of the dipolar orientations in the time window of approximately 20-30 ns can account for the experimental CCR. We compute the CCR of the dipole pair *H^N^ - N/C_α_ - H_α_* denoted by *R*_*d*(*H^N^-N*)/*d*(*C_α_-H_α_*)_ and defined as:

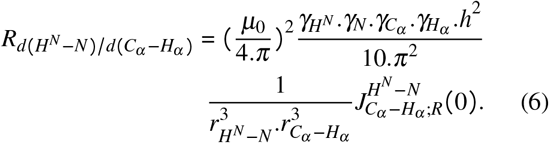

Here *γ_x_* denotes the gyromagnetic ratio and *r_x_* is the effective distance between nuclei *X*_1_ and *X*_2_. Here 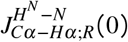 is the algebraic sum of zero frequency spectral function for both *H^N^ - N/C_α_ - H_α_* and *H^N^ - C_α_/N - H_α_* dipole pairs. This quantity has been computed numerically by integrating over the smoothened data, restricting up to 50 ns so that the initial decay of the correlations is entirely captured in the integration.

We divide the simulation trajectory after RMSD saturation into 8 equal windows, each consisting of configurations stored for 100 ns. This time interval is sufficient to guarantee the decay of the TDCFs so that each window can be treated as independent as far as the dipolar fluctuations are concerned. We compute the zero frequency spectral function for the dipolar fluctuations in each window given by the sum of the zero frequency spectral functions of both the direct and the cross dipolar pairs, as in the experiments. The theoretical CCR data is obtained from the spectral function for each window and finally averaged over the windows. The error in the mean is given by 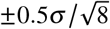 where *σ* being the standard deviation over the windows.

We consider in particular GB3 for which experimental data is available^44^. Fig. 3(c) shows the estimated theoretical CCR with the error bars along with available experimental data for GB3. We show linear regression through the data along with the upper and lower boundaries determined from the errors in the regression parameters. The slope and intercept obtained from linear regression are 0.23 and −0.46 respectively. The upper boundary around the linear regression line is plotted by considering minimum intercept and maximum slope. Similarly, lower boundary about the regression line is plotted considering maximum intercept and minimum slope. It turns out that the majority of observations fall within the boundaries, suggesting that the fluctuations up to 50 ns of the dipolar mutual orientation largely capture the experimental CCR. Some residues fall outside the lower limits: Ile 7, Asn8, Lys19, Val21, Asp22, Val39, Asp40. Val21 and Asp22 belongs to the loop region. The CCR rate is higher than the theoretical limit in particular for the residues adjacent to Glycine (Gly), for instance, Ile7, Asn8, Val39, and Asp40 having residue Gly9, Gly38, and Gly41 in their neighborhood respectively. Thus, typically, the dipolar fluctuations within the timescale of 50 ns well describe the experimental data for GB3.

*Correlated dipolar and dihedral fluctuations:* Good agreement between theoretical and experimental CCR for GB3 leads us to consider correlated dipolar and dihedral fluctuations for both the proteins. We compute the zero frequency spectral function for the dihedral fluctuations which we denote by *J_R_*(*θθ’*,0) for a given residue R over different windows as done for the dipolar fluctuations. We estimate the mean and the error in the mean considering data from all the windows.

The correlations between the zero frequency spectral functions of the dihedral and dipolar fluctuations for various residues are shown in Fig. 4. The theoretical error bars are also shown in the figure. Correlation is expressed in terms of weighted Pearson correlation with the inverse of the total error bar as weight. The weighted correlation formula for two variables x and y with the weight vector w, having the same length is given by;

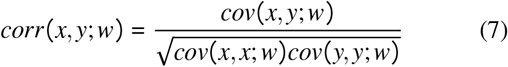

**FIG. 4.**
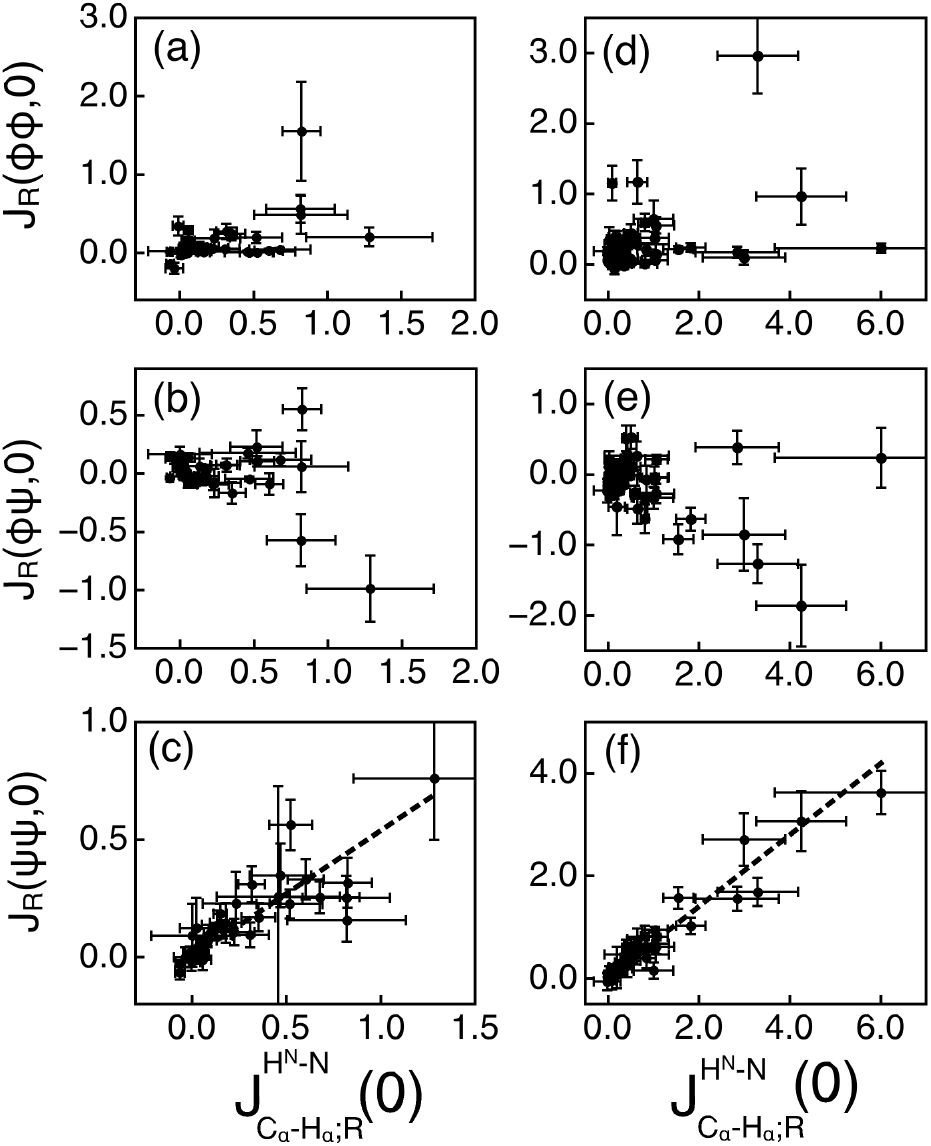
Correlation plot between zero frequency spectral functions of dihedral angle and dipole: (a) *J_R_*(*ϕϕ*,0), (b) *J_R_*(*ϕψ*,0), (c) *J_R_*(*ψψ*,0) with 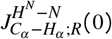 for GB3; (d) *J_R_*(*ϕϕ*,0), (e) *J_R_*(*ϕψ*,0), and (f) *J_R_*(*ψψ*,0) with 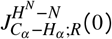 for Ub. Amberff is considered in both cases.

Where weighted covariance, *cov* (*x, y; w*) is given as,

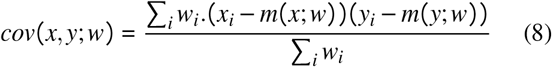

*m*(*x; w*) is the weighted mean. We take the weight of data points as the inverse of total error associated with each of them. Total error for any residue is considered as a simple algebraic sum of corresponding dihedral and bond fluctuation error.

Fig. 4(a)-(c) show data for GB3. The correlations are shown between 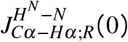 (Fig. 4(a)), *J_R_*(*ϕψ*,0) (Fig. 4(b)), and *J_R_*(*ψψ*,0)(Fig. 4(c)) respectively for GB3. While the correlation is poor or only moderate in case of 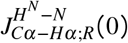 versus *J_R_*(*ϕϕ*,0) and 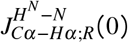 Versus *J_R_* (*ϕψ*, 0) data, it is strong for 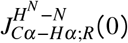 versus *J_R_* (*ψψ*, 0) data. The values of the weighted Pearson coefficients, denoted by 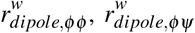 and 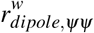 in Table.I confirm this trend. The data for Ub show a similar trend for 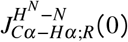 versus *J_R_*(*ϕϕ*,0), *J_R_*(*ϕψ*,0), and *J_R_*(*ψψ*,0) data shown in Fig.4(d)-(f). The weighted Pearson correlation coefficient values in Table.I also confirm this. We also show a linear regression plot in the case of 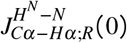 and *J_R_*(*ψψ*,0) data where the correlation is strong. Table. II lists the *R*^2^ values, slopes, and intercept obtained from linear regression for both proteins in the case of *ψ*.

**TABLE I.** Table of weighted Pearson correlation 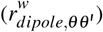

| Protein & Force field | $r_{dipole,\phi\phi}^w$ | $r_{dipole,\phi\psi}^w$ | $r_{dipole,\psi\psi}^w$ |
| --- | --- | --- | --- |
| GB3 with Amberff | 0.45 | -0.12 | 0.86 |
| Ub with Amberff | 0.45 | -0.39 | 0.94 |
| Ub with CHARMMff | 0.45 | -0.46 | 0.87 |

**TABLE II.** Table of linear regression (y = A*x + B)

| Protein & Force field | $R^2$ | A | B |
| --- | --- | --- | --- |
| GB3 with Amberff | 0.97 | 0.54 | 0.001 |
| Ub with Amberff | 0.99 | 0.7 | 0.008 |
| Ub with CHARMMff | 0.97 | 0.604 | 0.006 |

We check if the nature of correlation holds for another choice of force fields. We analyze data obtained for Ub using the CHARMM27 force field (For RMSD and *ϕ - ψ* plots, see SI Fig.S9 - Fig.S10). Figs. 5(a)-(c) show the correlation plots between zero frequency spectral function for the dipolar and dihedral fluctuations along with theoretical errors. The linear regression line is shown in the plot. The nature of the plots is similar to the results obtained for Ub from the Amber force field. The weighted Pearson correlation coefficients for the spectral functions (Table. I) are comparable with those obtained using the Amber force field. The correlation is stronger in case of 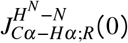 and *J_R_*(*ψψ*,0) data. The data for the linear regression data are listed in Table II. Fig 5(d) shows correlation plot of *τ_R,H^N^-N/C_α_-H_α__* and *τ_R,ϕϕ_* for dihedral angle *ϕ* considering all three cases. The correlation plot suggests that *τ_R,ϕϕ_* is poorly correlated with *τ_R,H^N^-N/C_α_-H_α__* for all the cases (Fig. 5(d)). Fig 5(e) shows a moderate correlation between *τ_R,ϕψ_* and *τ_R,H^N^-N/C_α_-H_α__*· *τ_R,ψψ_* is strongly correlated with *τ_R,H^N^-N/C_α_-H_α__* (Fig 5(f)). We denote Pearson correlations by *r_dipole, ϕ_* for correlation between dipolar orientation and *ϕ* and that between dipolar orientation and *ψ* by *r_dipole, ψ_*. We plot the histograms of the correlation coefficients *H*(*r_dipole,ϕ_*) for different cases in Fig. 6(a). The histograms have a peak near zero value suggesting that *ϕ* fluctuations are largely uncorrelated to the dipolar fluctuations. Histograms *H*(*r_dipole,ψ_*) Fig. 6(b), on the other hand, show strong correlations for a large number of residues for all three cases. Thus, our data suggest common a feature of microscopic motions in these proteins.

**FIG. 5.**
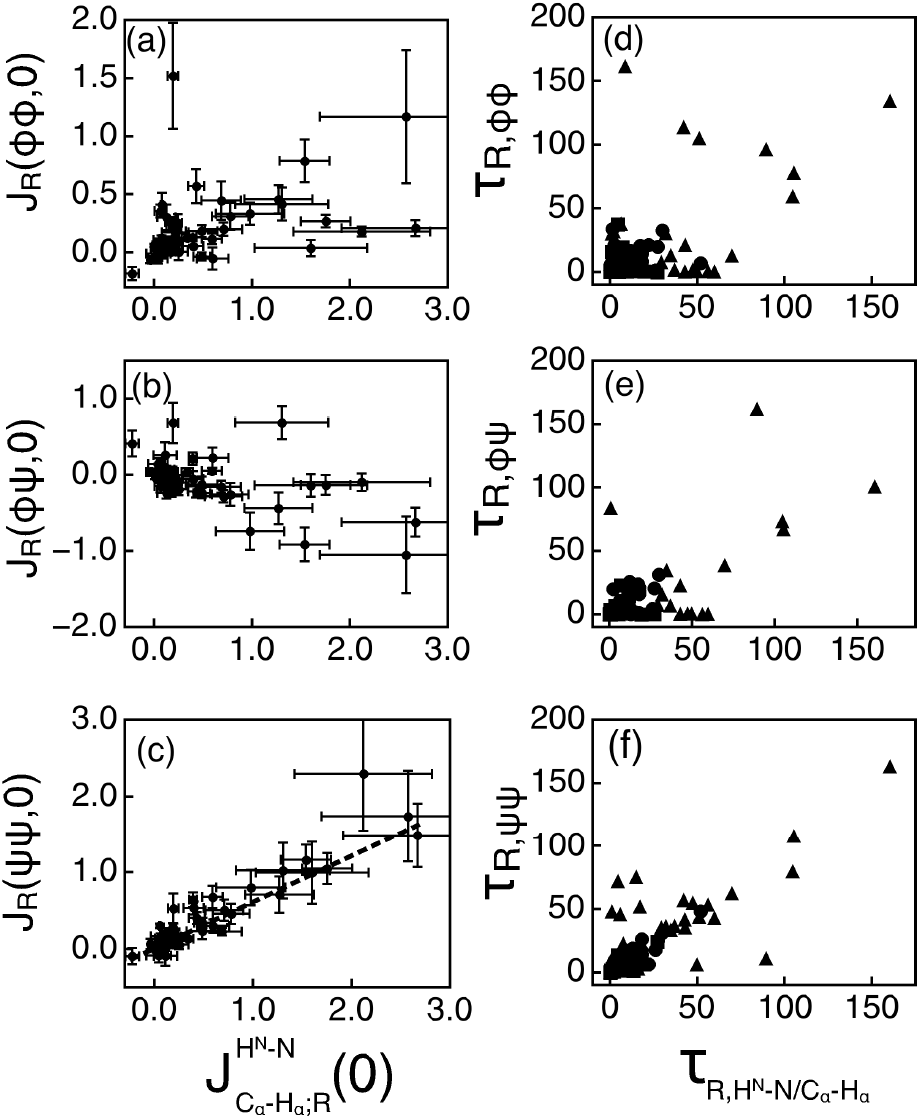
Correlation plot between zero frequency spectral function of dihedral angle and dipole: (a) *J_R_*(*ϕϕ*,0), (b) *J_R_*(*ϕψ*,0), (c) *J_R_*(*ψψ*,0) with 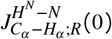 for Ub, considering CHARMM ff. Correlation plot of timescales of dihedral and dipolar fluctuations, (d) *τ_R,ϕϕ_* (e) *τ_R,ϕψ_* and (f) *τ_R,ψψ_* with timescales of dipole *H^N^ - N/C_α_ - H_α_*, (*τ_R, H^N^-N/C_α_-H_α__*). Different cases are shown in different symbols: ■ for GB3 using Amberff, ● for Ub using Amberff and ▲ for Ub using CHARMMff.

**FIG. 6.**
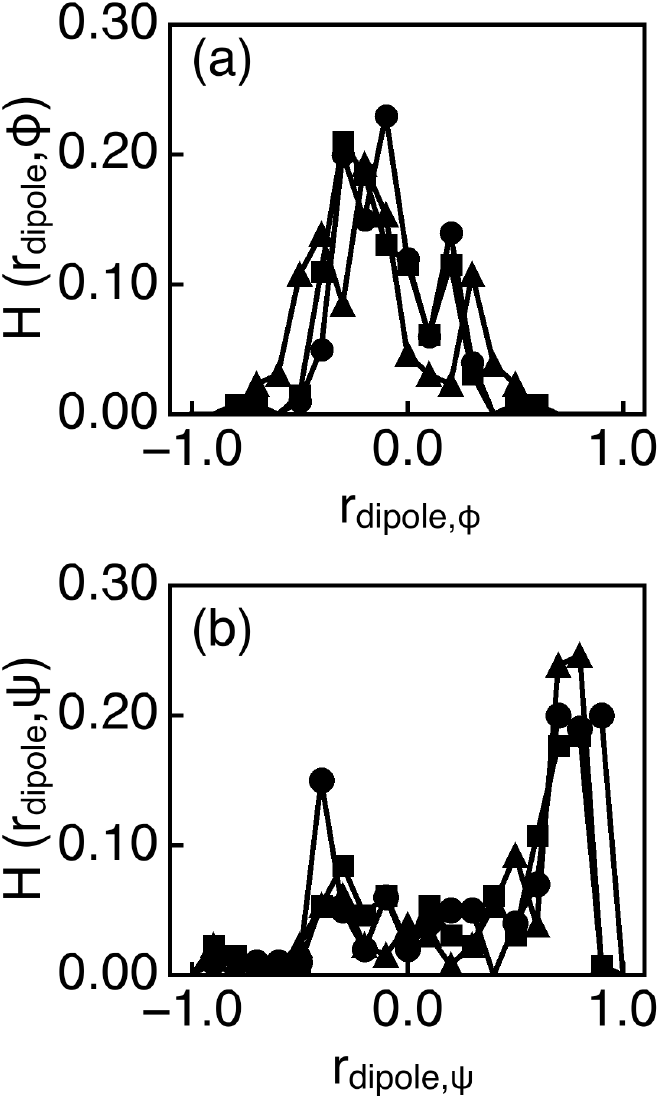
(a) Histogram *H*(*r_dipole, ϕ_*) of Pearson correlation coefficient values for fluctuation of dipole vector with dihedral *ϕ*. (b) Histogram *H*(*r_dipole, ψ_*) of correlation coefficient values for fluctuation of dipole vector with dihedral *ψ*. Different cases are shown in different symbols. ● for GB3 using Amberff, ■ for Ub using Amberff and ▲ for Ub using CHARMMff.

## IV. CONCLUSIONS

To summarize we show that fluctuations at the timescale of a few tens of nanoseconds can capture the experimental CCR of dipolar fluctuations in a protein. Within this timescale, the zero frequency spectral functions of the dipolar fluctuations and intra-residual backbone dihedral *ψ* show good correlations. The correlation nature is independent of the protein and force fields. Our study throws light to universal feature of backbone motions formed by the peptide bonds which merits further investigations.

## Supporting information

Suppplementary File

## ACKNOWLEDGMENTS

A.G.M acknowledges DST India, for an Inspire Fellowship (IF170961). We acknowledge Rahul Karmakar for critically reading the manuscript.

