## Supplementary material for "Correlated dipolar and dihedral fluctuations in a protein": Suppplementary File

January 28, 2021

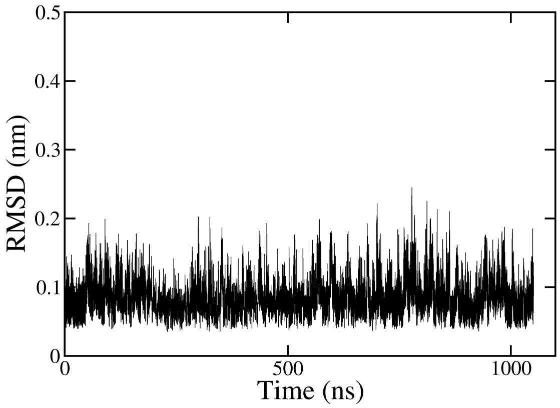


Supplementary Figure, Figure S1: RMSD plot of GB3 over 1.05 $\mu$s using Amber force field.

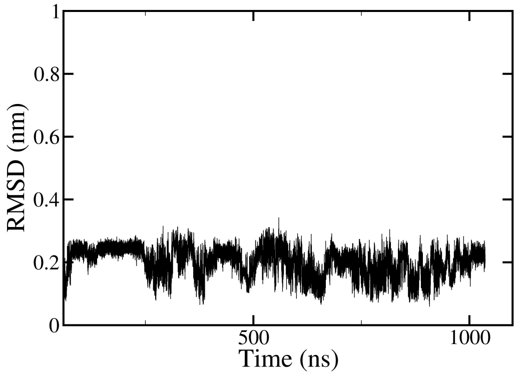


Supplementary Figure, Figure S2: RMSD plot of Ub over 1.05 $\mu$s considering Amber force field.

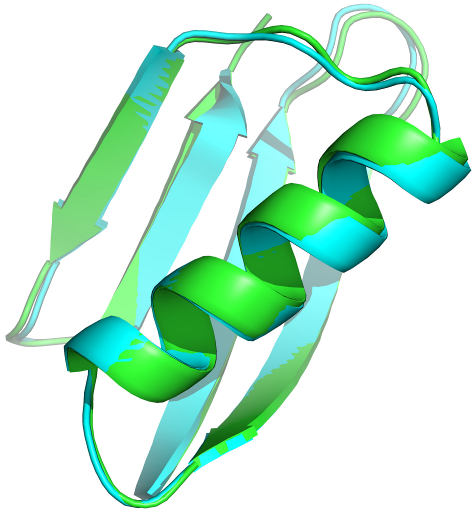

Supplementary Figure, Figure S3: Overlapped image of initial and average structure of GB3. Green color represent initial structure and cyan represents average structure.

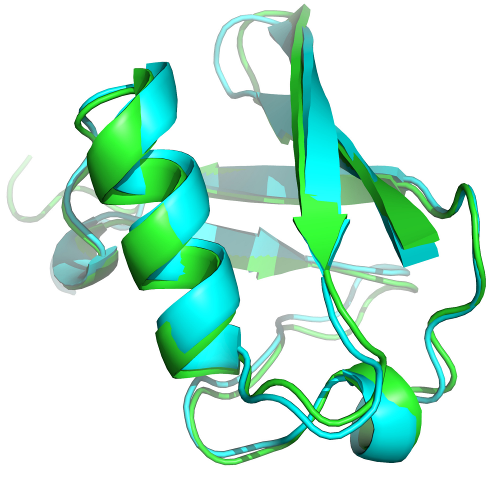

Supplementary Figure, Figure S4: Overlapped image of initial and average structure of Ub. Green color represent initial structure and cyan represents average structure.

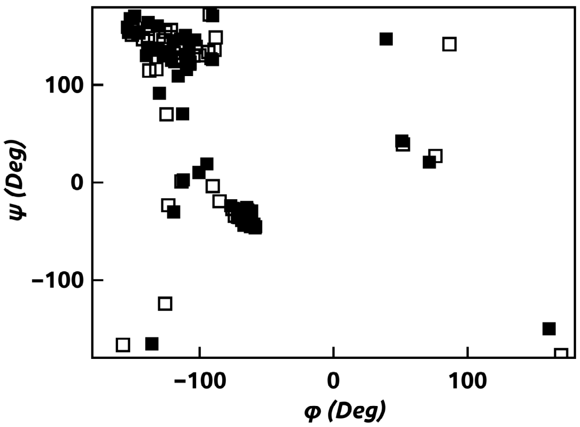


Supplementary Figure, Figure S5: $\psi - \phi$ Correlation plot of residues in GB3 using Amber force field; Filled rectangle represents the crystal structure and hollow rectangle shows simulated average structure obtained from the equilibrated trajectory

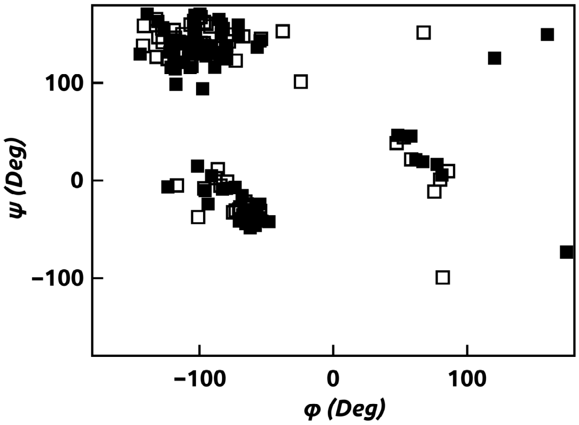


Supplementary Figure, Figure S6: $\psi - \phi$ Correlation plot of residues in Ub using Amber force field; Filled rectangle represents the crystal structure and hollow rectangle shows simulated average structure obtained from the equilibrated trajectory

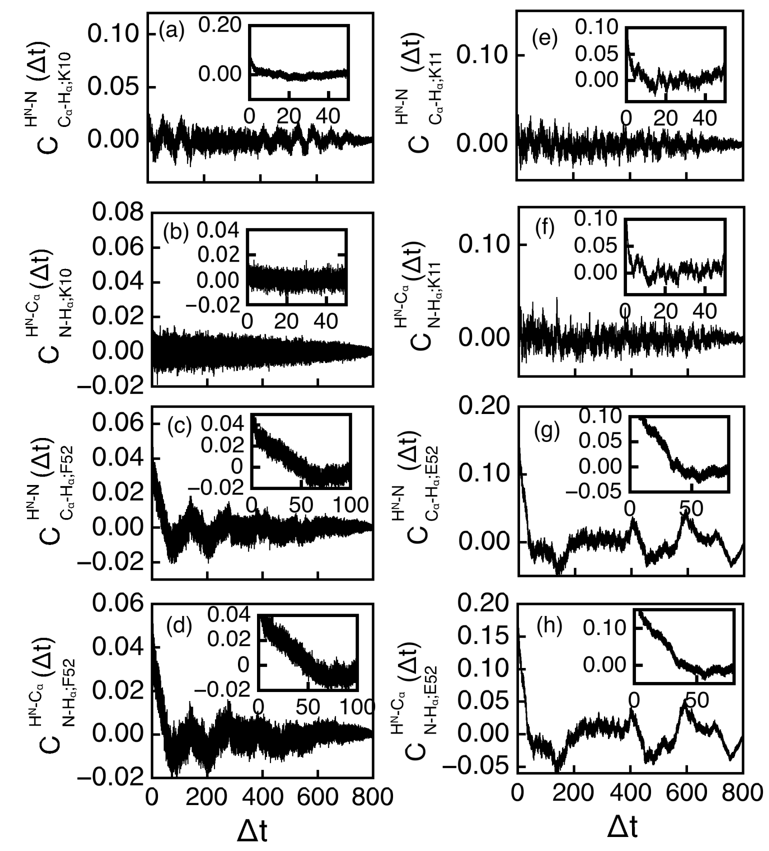

Supplementary Figure, Figure S7: TDCF of dipolar fluctuation. Residue Lysine (K10) (a-b), Phenylalanine (F52) (c-d) of GB3 protein and residue Lysine (K11) (e-f), Glutamic acid (E52) (g-h) of Ub protein. Inset shows short time behaviour. For each residue both $H^N - N/C_\alpha - H_\alpha$ and $H^N - C_\alpha/N - H_\alpha$ dipoles are considered.

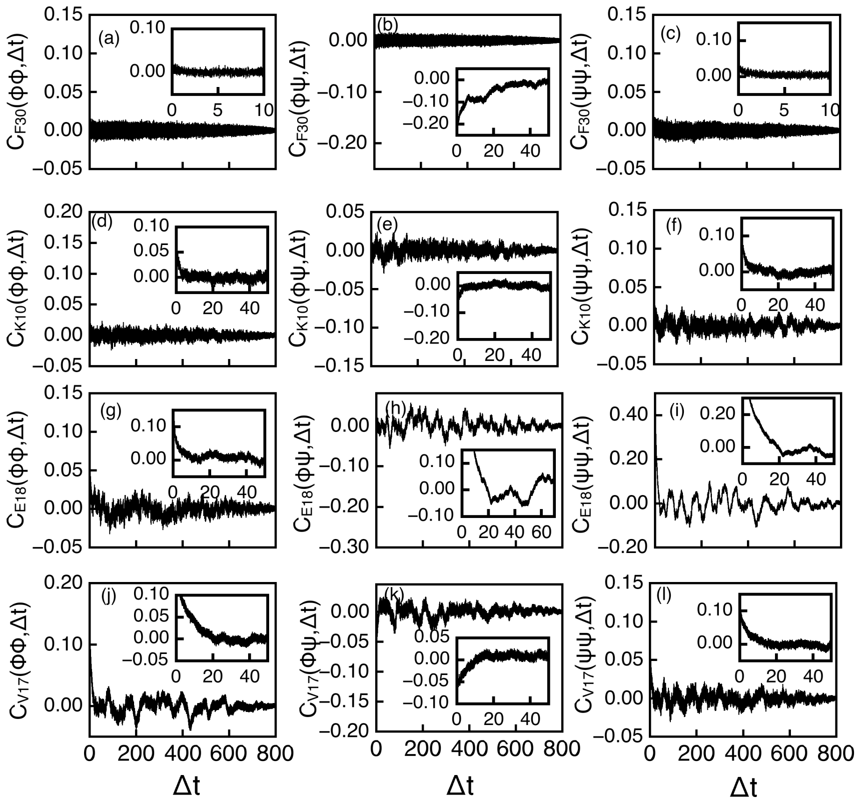

Supplementary Figure, Figure S8: TDCF of dihedral fluctuations. For GB3, Phenylalanine (F30) (a-c) and Lysine (K10) (d-f) are considered. Glutamic acid (E18) (g-h) and Valine (V17) (i-k) are considered for Ub. Inset shows short time behaviour.

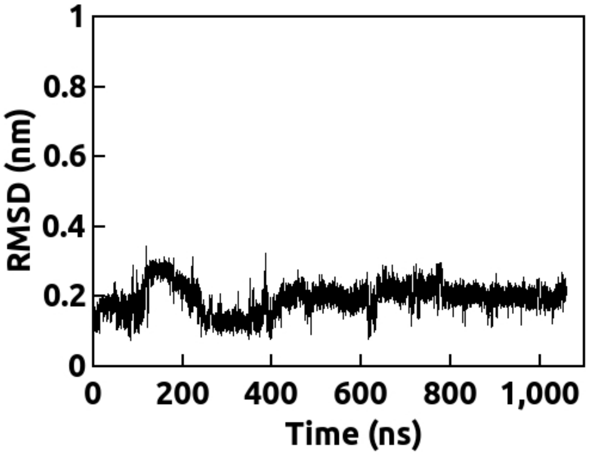


Supplementary Figure, Figure S9: RMSD plot of ubiquitin over 1.05 $\mu$s using CHARMM forcefield.

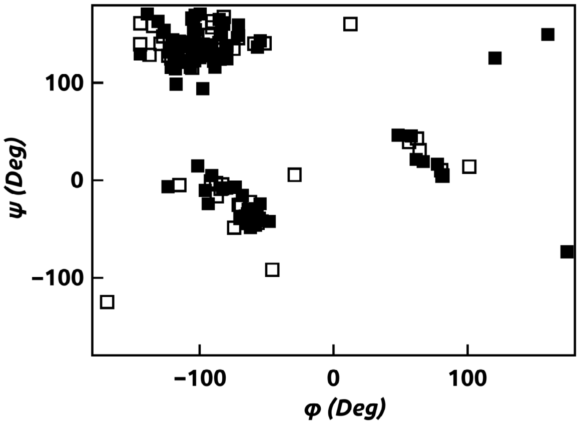


Supplementary Figure, Figure S10: $\psi - \phi$ Correlation plot of residues in Ub using CHARMM field; Filled rectangle represents the crystal structure and hollow rectangle shows simulated average structure obtained from the equilibrated trajectory.
